# I-Impute: a self-consistent method to impute single cell RNA sequencing data

**DOI:** 10.1101/772723

**Authors:** Xikang Feng, Lingxi Chen, Zishuai Wang, Shuai Cheng Li

**Author notes:** Equal contributor (Xikang Feng, Lingxi Chen).

## Abstract

Single-cell RNA-sequencing (scRNA-seq) is essential for the study of cell-specific transcriptome landscapes. The scRNA-seq techniques capture merely a small fraction of the gene due to “dropout” events. When analyzing with scRNA-seq data, the dropout events receive intensive attentions. Imputation tools are proposed to estimate the values of the dropout events and de-noise the data. To evaluate the imputation tools, researchers have developed different clustering criteria by incorporating the ground-truth cell subgroup labels. There lack measurements without cell subgroup knowledge. A reliable imputation tool should follow the “self-consistency” principle; that is, the tool reports the results only if it finds no further errors or dropouts from the data. Here, we propose “self-consistency” as an explicit evaluation criterion; also, we propose I-Impute, a “self-consistent” method, to impute scRNA-seq data. I-Impute leverages continuous similarities and dropout probabilities and refines the data iteratively to make the final output self-consistent. On the *in silico* data sets, I-Impute exhibited the highest Pearson correlations for different dropout rates consistently compared with the state-of-art methods SAVER and scImpute. On the datasets of 90.87%, 70.98% and 56.65% zero rates, I-Impute exhibited the correlations as 0.78, 0.90, and 0.94, respectively, between ground truth entries and predicted values, while SAVER exhibited the correlations as 0.58, 0.79 and 0.88, respectively and scImpute exhibited correlations as 0.65, 0.86, and 0.93, respectively. Furthermore, we collected three wetlab datasets, mouse bladder cells dataset, embryonic stem cells dataset, and aortic leukocyte cells dataset, to evaluate the tools. I-Impute exhibited feasible cell subpopulation discovery efficacy on all the three datasets. It achieves the highest clustering accuracy compared with SAVER and scImpute; that is, I-Impute displayed the adjusted Rand indices of the three datasets as 0.61, 0.7, 0.52, which improved the indices of SAVER by 0.01 to 0.17, and improved the indices of scImpute by 0.19 to 0.4. Also, I-impute promoted normalized mutual information of the three datasets by 0.01 to 0.09 comparing with SAVER, and by 0.15 to 0.34 comparing with scImpute. I-Impute exhibits robust imputation ability and follows the “self-consistency” principle. It offers perspicacity to uncover the underlying cell subtypes in real scRNA-Seq data. Source code of I-Impute can be accessed at **https://github.com/xikanfeng2/I-Impute.**

## Background

Single-cell RNA-sequencing (scRNA-seq) is becoming essential for the study of cell-specific transcriptome landscapes (1). It demonstrates robust efficacy in capturing transcriptome-wide cell-to-cell heterogeneity with high resolution (2–5). With meta information such as time series or patient histology, scRNA-seq has the potential to decipher the underlying patterns in cell cycles (6–8), complex diseases (9–11), and cancers (8, 12–16).

A count matrix captures expression profiles with genes as rows and cells as columns, and the measurements of count as the matrix entries. scRNA-seq captures a small fraction of the gene due to “dropout” events; that is, scRNA-seq produces zero-inflated count matrices, only about 10% entries are non-zero values (17). Usually, the dropout events occur as the truly expressed transcripts may be missed during sequencing in some cells, and the dropout rate is protocol-dependent (18). When analyzing with scRNA-seq data, the excess zero counts brought by dropout issue are worth for attention. Otherwise, the zero count distribution from different protocols may lead diverging potency, which may weaken the credibility of the downstream analysis results (18).

The conventional downstream analyses include clustering, cell type recognition, dimension reduction, differential gene expression analysis, identification of cell specific genes and reconstruction of differentiation trajectory on zero-inflated single-cell gene expression data (18). The credibility of the aspects mentioned above depends on the exactitude of expression profiling; therefore, it is vital to amend the false zeros induced by dropout events. In past years, some scRNA-seq methods choose to conduct clustering, cell type recognition, and dimension reduction tasks by implicitly incorporating dropout events (19–22). While at this moment imputation before downstream analysis becomes mainstream, scRNA-seq imputation tools have rapidly emerged to tackle the rampant dropouts. SAVER (23) imputes by borrowing information across genes and using the Bayesian approach to estimate the expression levels. It aims to reduce meaningless biological variation and retain valuable biological variation. While SAVER would unfairly adjust all gene expression levels including the actual non-expression of genes, hence possibly interject new biases and abolish real biological meanings. scImpute (18) is designed to identify dropout values with Gamma-Normal mixture model firstly, and then do imputation on dropout events by borrowing information from similar cells, with the expression level of un-dropout events unchanged. It automatically excludes the outlier cells and their gene information, which are likely to influence the original imputation values. While scImpute is not good at the extremely sparse datasets.

On *in silico* data where the ground truth counts are known, the root mean square error (RMSE) between imputed and ground truth entries is the most common metrics for imputation evaluation (24). For wetlab data sets, the ground truth counts for missing events are unknown. Researchers randomly remove non-zeros entries and employ the imputation tools to impute these removed entries. Then they calculated the RMSE for removed entries as a criterion to evaluate imputation tools (24, 25). Also, researchers tend to implicitly validate imputation efficacy by checking whether imputed data promotes the downstream analysis result or not. For instance, clustering measurements such as adjusted Rand index (ARI), normalized mutual information (NMI), silhouette width (SW), and within-cluster sum of squares are commonly adopted for scRNA-seq imputation evaluation (18, 26). Nevertheless, the aforementioned clustering measurements require the true cluster labels. There lack explicit measurements for wet-lab data imputation.

A reliable imputation tool should assume its output contains no dropout or errors; that is, if we feed the output to the imputation tool again by eliminating a certain amount of entries, the tool should be able to reproduce these entries. We refer this property as the “self-consistency” principle for imputation. Therefore, in this study, we propose the concept of “self-consistency” as a criterion to evaluate the reliability of imputation tools by just using the input count matrix itself. We measured the self-consistency of the state of the art imputation tools, and as well as developed a self-consistent method “I-Impute” for scRNA-seq data imputation. On the *in silico* data sets, I-Impute exhibited consistently the highest Pearson correlations for different dropout rates compared with the state-of-art methods SAVER and scImpute. On the datasets of 90.87%, 70.98% and 56.65% zero rates, I-Impute exhibited the correlations as 0.78, 0.90, and 0.94, respectively, between ground truth entries and predicted values, while SAVER exhibited the correlations as 0.58, 0.79 and 0.88, respectively and scImpute exhibited correlations as 0.65, 0.86, and 0.93, respectively. Furthermore, several discrete cell subpopulations have been reported in scRNA-Seq data collected from the wet lab; the identification of subpopulations of cells is crucial (27). Here, we collected three wetlab datasets, mouse bladder cells dataset, embryonic stem (ES) cells dataset, and aortic leukocyte cells dataset to evaluate the tools. I-Impute exhibited feasible cell subpopulation discovery efficacy on all the three datasets. It achieves the highest clustering accuracy compared with SAVER and scImpute; that is, I-Impute displayed the adjusted Rand indices of the three datasets as 0.6054, 0.7047, 0.5220; while SAVER and scImpute have the indices as 0.5253 and 0.1937, 0.6920 and 0.3574, 0.3605 and 0.3377, respectively. Also, I-impute promoted normalized mutual information of the three datasets from 0.7085 and 0.45 to 0.7892, from 0.7329 and 0.5258 to 0.7444, from 0.6837 and 0.6237 to 0.7728, respectively.

## Results

### Evaluating the self-consistency of existing imputation tools in synthetic data

To evaluate the imputation tools, we applied the R package Splatter (28) to generate scRNA-seq reads count data. We simulated 150 cells of three groups, each with 2,000 genes. Then we generated three sparse matrices by setting the dropout rates as 88.45%, 63.29%, and 45.16%; and their corresponding zero rates are 90.87%, 70.98%, and 56.65%, respectively.

Above all, we validated whether existing imputation tools were self-consistent. We call a method *f* : *x* →*x* is self-consistent if the root mean square error (RMSE) between *x*_*output*_ and *f*(*x*_*output*_) is less than prespecified thresholds *θ*, where *x*_*output*_ = *f*(*x*_*input*_). Considering the imputation process as a complex function *f*_*imputation*_ that maps the zero-inflated matrix into an output matrix of the same shape. A reliable function *f*_*imputation*_ should procure the recovered matrix with no noise nor missing entries, aka ‖ *x*_*output*_ − *f*_*imputation*_(*x*_*output*_) ‖2 ≤*θ*. As illustrated in **Table 1**, assume the cut-off value *θ* is 0.1, SAVER and sc Impute is self-inconsistent. scImpute exhibits self-inconsistency with RMSE value of 7.346 in 88.45% dropout data, of 0.2392 in 63.29% dropout data, of 0.2677 in 45.16% dropout data. SAVER reveals self-inconsistency with RMSE value of 0.5613, 1.0245, and 1.3561 in above three datasets, respectively. Nevertheless, incorporating ground truth group labels, traditional evaluation metrics represented that SAVER out-performed scImpute in respect to adjusted Rand index (ARI), normalized mutual information (NMI), and silhouette width (SW) (see **Table S1**).

**Table 1.** Self-consistency on synthetic data. NA denotes not applicable.

|  | SAVER | scImpute | I-Impute |
| --- | --- | --- | --- |
| 88.45% dropout | 0.5613 | 7.3460 | 0.0936 |
| Self-consistent (<0.1) | x | x | ✓ |
| 63.29% dropout | 1.0245 | 0.2392 | 0.0806 |
| Self-consistent (<0.1) | x | x | ✓ |
| 45.16% dropout | 1.3561 | 0.2677 | 0.0381 |
| Self-consistent (<0.1) | x | x | ✓ |

Therefore, in this study, we build a two-pronged method “I-Impute” which satisfies both self-consistency principle and as well as existing imputation metrics (ARI, NMI, and SW). As illustrated in **Figure 1A**, we first developed “C-Impute by adopting continuous similarities and dropout probabilities to infer the missing entries. Then, I-Impute invokes SAVER as a subrourtine to preprocess the data, and then employ C-Impute iteratively to processed data (see **Figure 1B**). With *n* times of iterations, the final imputed result turns to self-consistent, with RMSE value of 0.0936, 0.0806, and 0.0381 in three synthetic datasets, respectively (see **Table 1**).

**Fig. 1.**
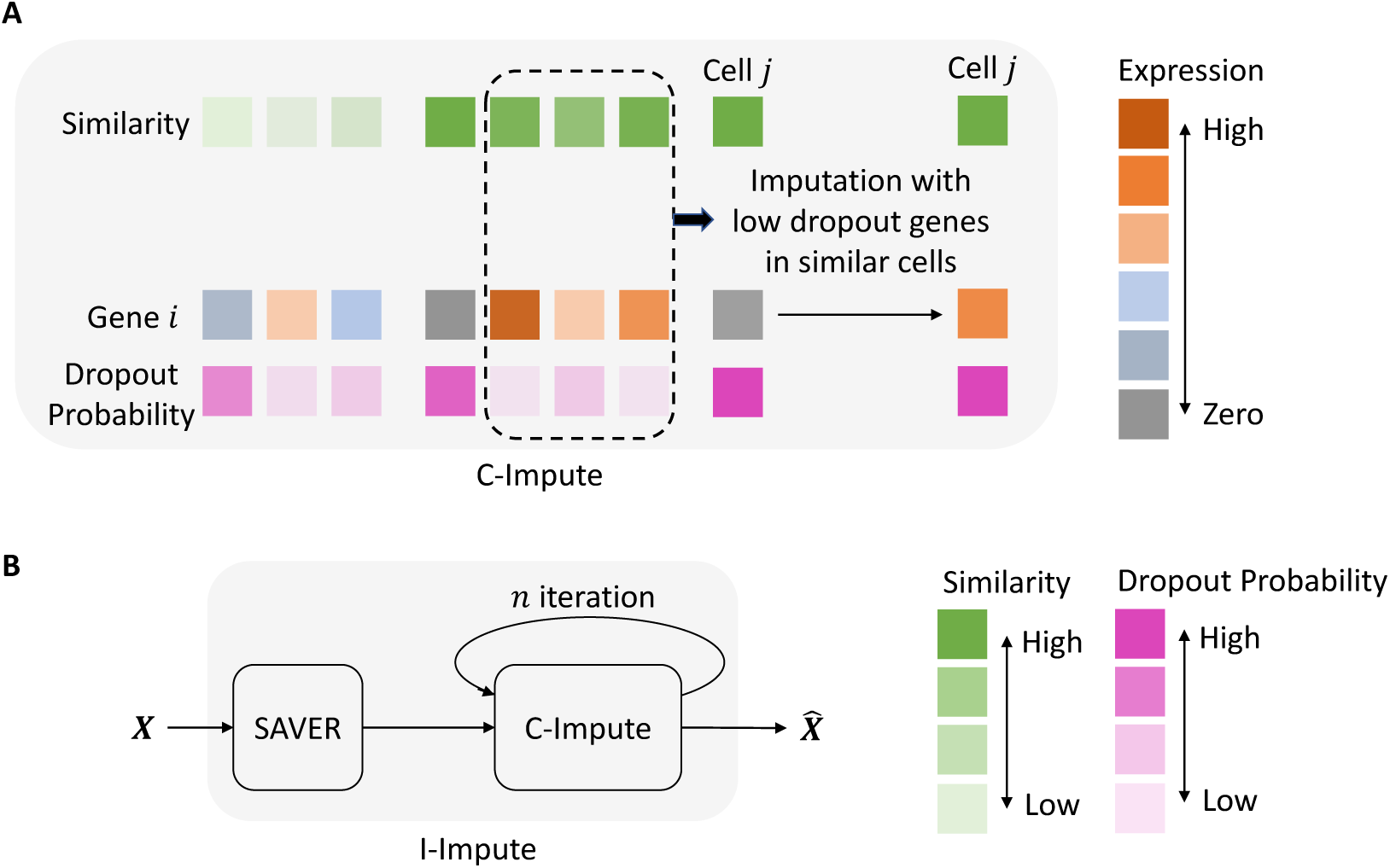
Illustration of I-Impute architecture.

### I-Impute recovers gene expression affected by dropouts in synthetic data

To validate the performance of I-Impute, we painted the heatmap of the raw matrix, 88.45% dropout matrix, and recovered matrices, respectively (see **Figure 2A-F**). I-Impute recovered the most similar pattern than SAVER, scImpute, and C-Impute. As illustrated in **Figure 2G**, SAVER pulled down a large part of entries from their raw, leading to the lowest Pearson correlation 0.58 between ground truth and prediction. scImpute and C-Impute casted some high expressed entries into zero, which introduces new bias after imputation (see **Figure 2H-I**). With no extreme pull-down or pull-up prediction, I-Impute exhibited the most robust recovery potency with the highest Pearson correlation 0.78 (see **Figure 2J**). In terms of 63.29% and 45.16% dropout rate, I-Impute also manifested the highest Pearson correlation of 0.90 and 0.94, respectively (see **Table S4**).

**Fig. 2.**
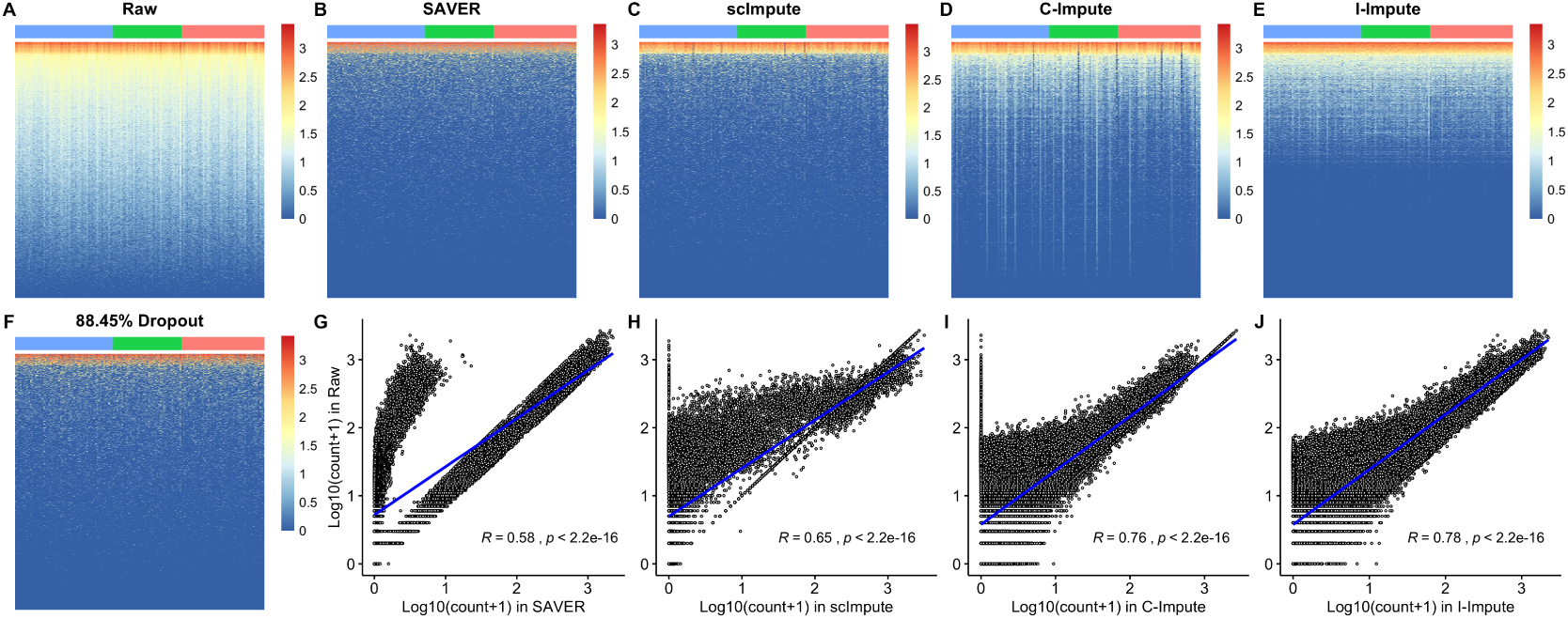
Imputation performance on synthetic data (A-F) Heatmap plots. Blue, green, and red tiles represent different cell groups (C-J) Scatter plots, Pearson correlation between ground-truth entries and imputed values are calculated.

The t-SNE embedding plots of the raw matrix, 88.45% dropout matrix, and recovered matrices demonstrate that SAVER, C-Impute, and I-Impute recover the missing entries while preserving cell subgroups structures well (see **Figure 3A-F**). Silhouette width (SW) further validated that the in-group similarity and out-group separation were enhanced after the imputation by SAVER, C-Impute, and I-Impute. That is, the average silhouette value increased from 0.0862 (dropout data) to 0.1075 (SAVER), 0.1705 (C-Impute), and 0.2429 (I-Impute), respectively (see **Table S1**). **Figure 3G** demonstrates that I-Impute achieves the most noticeable improvement, while scImpute illustrates lower SW values than dropout data. Next, we applied hierarchical clustering into all matrices, and adopted the adjusted Rand index (ARI) and normalized mutual information (NMI) as metrics of clustering accuracy. ARI and NMI measure the overlap between the inferred groups and ground-truth clusters; a score of 0 implies random labelling and 1 indicates perfect inference. In **Figure 3G**, I-Impute outperforms all four tools and exhibits the best subpopulation identification strength, with the highest clustering accuracy (ARI: 0.8721, NMI: 0.8521, see **Table S1**). Experiments on 63.2% and 45.16% dropout rate data sets also proved that I-Impute produced the best recovered matrices; with ARI 1.0, NMI 1.0, SW 0.3908 for 63.2% dropout rate, and ARI 0.9801, NMI 0.9710, and SW 0.4123 for 45.16% dropout rate (see **Table S2-S3**).

**Fig. 3.**
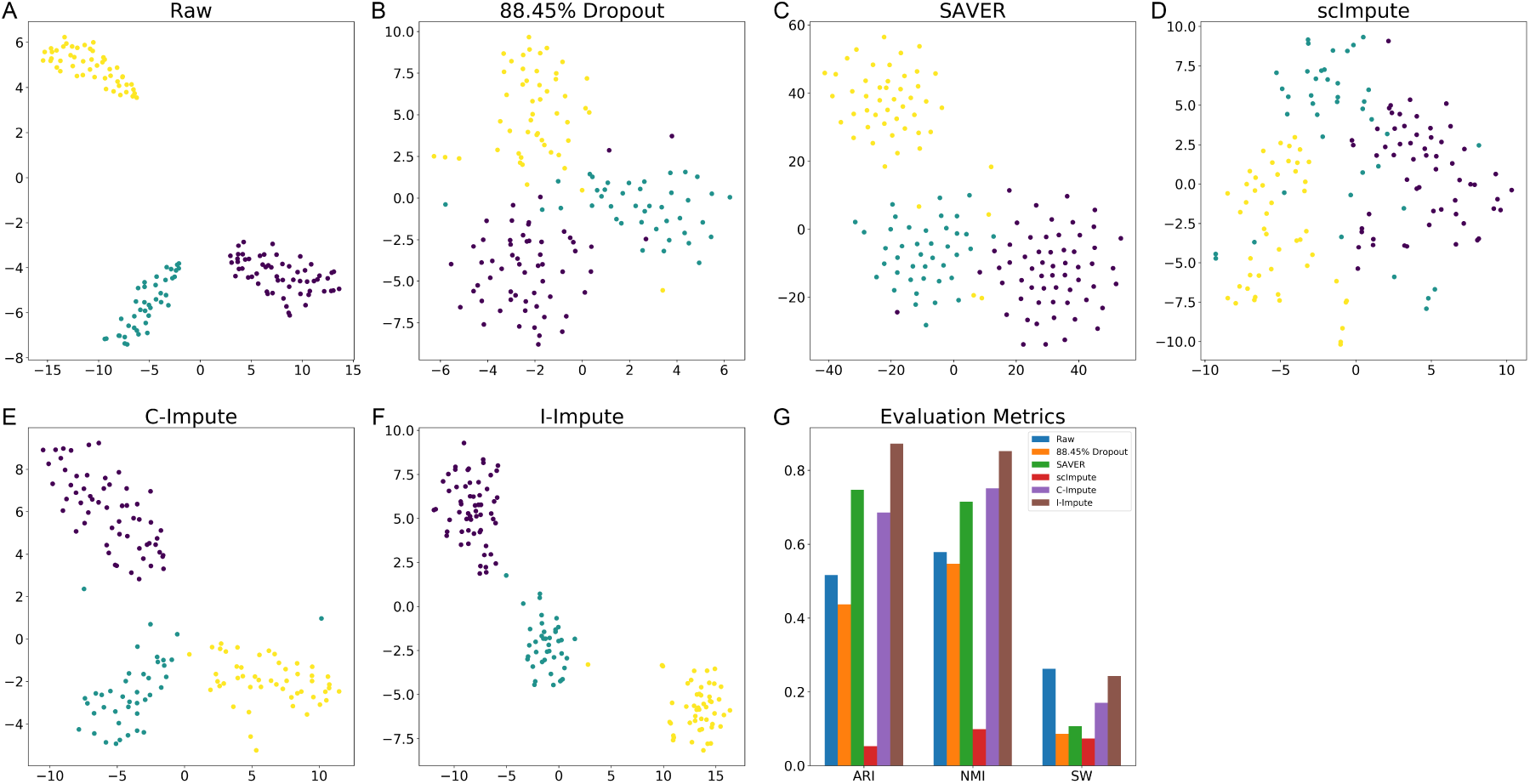
Imputation performance on synthetic data (A-F) t-SNE plots (G) evaluation metrics.

Overall, the synthetic experiment demonstrates that by incorporating C-Impute to refine the SAVER processed data iteratively, I-Impute outcomes the SAVER’s inconsistency and enhances the imputation potency.

### I-Impute promotes cell subpopulation identification in real data sets

To manifest how I-Impute can help to identify cell subpopulations, we utilized three real scRNA-Seq datasets as the benchmark. The first one is a dataset of mouse Bladder cells which contains 162 cells of three cell types. Due to dropout events, 73.5% of read counts in the raw count matrix are zeros. We evaluate the imputation power by reviewing the tSNE embedding result and silhouette width (SW). ScImpute mixs part of Unknown-type cells (the purple dots) with the Fibroblasts-1 cells (the blue dots) and Fibroblasts-2 cells (the yellow dots). While SAVER, C-Impute and I-Impute distinguish the Unknown-type cells from Fibroblasts-1 cells and Fibroblasts-2 cells well. Overall, compared with raw and other imputed data, the I-Impute produce the most compact clusters with highest silhouette width of 0.1758 (**Figure 4A**). We then compare the hierarchical clustering accuracy, ARI and NMI. All the two measurements intimate that with 0.6054 ARI and 0.7892 NMI, I-Impute heads to the best clustering effect as compared with raw (ARI:0.1937, NMI:0.45) and the imputation by SAVER (ARI:0.5253, NMI:0.7085), scImpute (ARI:0.1937, NMI:0.45), or C-Impute (ARI:0.1664, NMI:0.4317) (**Figure 4A**, **Table S5**).

**Fig. 4.**
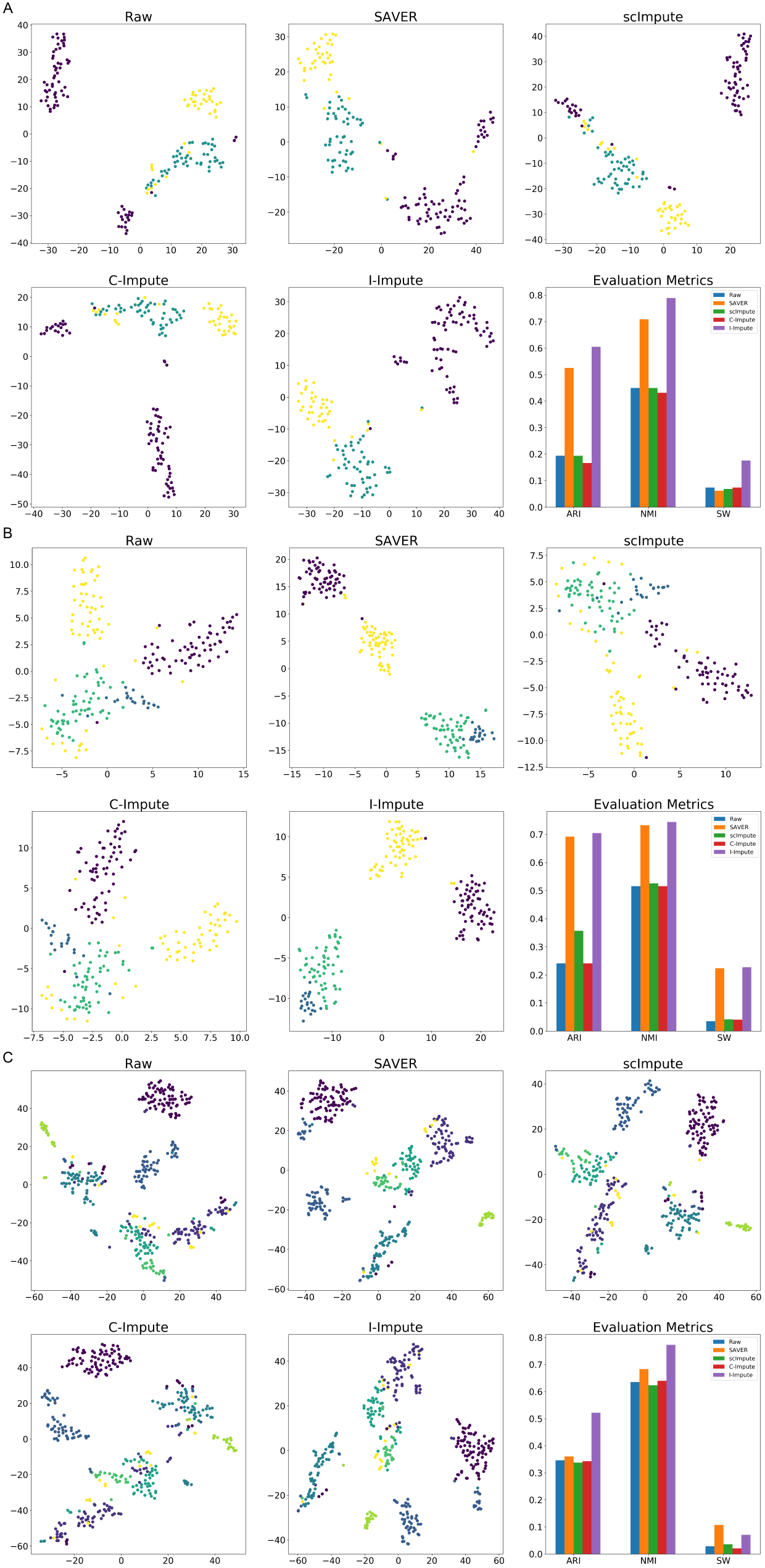
Imputation performance on real datasets (A-C) t-SNE plots and evaluation metrics for mouse bladder cells, embryonicstem cells, and aortic leukocyte cells, respectively

We also adopt I-Impute to a mouse embryonic stem (ES) cells dataset. This dataset contains 2717 cells of four cell types (mouse ES cells sample 1, mouse ES cells LIF 2 days, mouse ES cells LIF 4 days and mouse ES cells LIF 7 days). Due to the running time limitation of scImpute for large cells dataset, we randomly selected 200 cells and no subpopulations and genes were excluded during this process. Due to dropout events, 67.0% of read counts in the raw count matrix are zeros. Overall, **Figure 4B** illustrate that SAVER and I-Impute achieved the unprecedented imputation power than other tools. Given by the 2D t-SNE embedding space, SAVER and I-Impute separate 2 days cells (the yellow dot) from 4 days cells (the green dots) and 7 days cells (the blue dots) well. The evaluation of Silhouette width, adjusted Rand index, and normalized mutual information further demonstrated that I-Impute (ARI:0.7047, NMI:0.7444, SW:0.2275) produced a more tight and accurate in-cluster structure than SAVER (ARI:0.692, NMI:0.7329, SW:0.2235)(**Table S6**). This evidence shows the strong ability of I-Impute to identify cell subpopulations despite 67.0% missing rate.

Finally, we used the mouse Aortic Leukocyte cells dataset to test I-Impute. This dataset contains 378 cells of six cell types (B cells, T cells, T memory cells, Macrophages, Nuocytes, and Neutrophils). Due to dropout events, 91.2% of read counts in the raw count matrix are zeros. SAVER and I-Impute grouped the T memory cells (the yellow dots) into big cluster, while in raw data and other imputed matrices, T memory cells are separated into different clusters (see **Figure 4C**). Even though I-Impute with silhouette width of 0.0711 dose not recover a more compact structure than SAVER, it outperforms all other tools in hierarchical clustering tasks with highest ARI (0.522) and NMI (0.7728) (**Table S7**). Overall, I-Impute predicts missing values in scRNA-seq data and improves the discovery of cell subpopulations.

## Methods

### C-Impute

We propose “C-Impute” by adopting a flexible continuous similarity and Lasso penalty in objective function (see **Figure 1A**).

### Data prepossessing

The input of the method is a count matrix 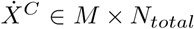 which contains rows as genes and columns as cells, where *M* and *N*_*total*_ represent the total number of genes and cells correspondingly. The dropout values are replaced by zero counts.

Then, normalization, dimension reduction, and outlier removal are conducted the same as in scImpute (18). Finally, we obtained outlier removed matrix *X ∈ M* ×*N* and *Z ∈ K* ×*N*, where *K* is the reduced dimensionality of metagenes, *N* is the number of remained cells.

### Affinity matrix constructing

Based on the outlier removed ***Z*** matrix, cell affinity matrix *A ∈ N* ×*N* can be computed with Euclidean distance and Gaussian Kernel:

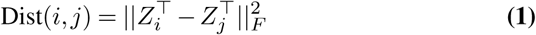

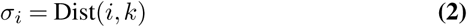

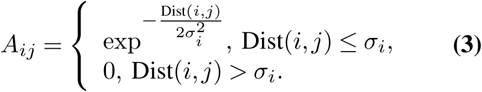

Where *i, j* represent two different cell indices, 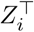 and 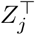 indicate the principle components of *i*-th and *j*-th cell respectively. For *i*-th cell, the kernel width will be set to the distance between it and its *n*-nearest neighbor, cell *k*, which stands for the cell whose distance to cell *i* is *n*-th smallest in all other cells, where *n* is a hyper-parameter.

### Identification of dropout values and calculating dropout rate

With transformed gene expression matrix *X*, we utilize a statistical model to infer which entries are influenced by the dropout effects. Instead of treating all zero values as missing entries, we use the Gamma-Normal mixture model to learn whether a zero observation originates from dropout or not. We use the Normal distribution to present the actual gene expression level and Gamma distribution to take into account for dropout events. Since the transformed matrix X contains no longer integers, we cannot adopt zero-inflated negative binomial (ZINB) distribution.

For *i*-th gene and its observed value *x* in prepossessed gene profiling *X*_*i*_, the Gamma-Normal mixture model will be:

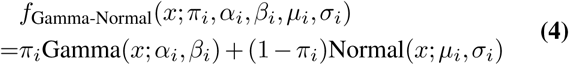

Where *π*_*i*_ is the dropout rate of gene *i, α*_*i*_ and *β*_*i*_ is the shape and rate parameter of Gamma distribution respectively, *µ*_*i*_ and *σ*_*i*_ are the mean and standard deviation of Normal distribution. The estimated model parameters 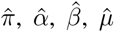 and 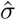 are obtained by Expectation-Maximization (EM) algorithm. Then, we can calculate the dropout probability matrix *D ∈ M* × *N*.

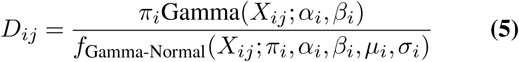

The purpose of this mixture model is to identify the observed value is a dropout value or not, since zero can be caused by a technical error or may the actual value of its expression. If a gene has high expression and low variation in most its similar cells, a zero count will have high dropout probability and more likely to be a dropout value; otherwise, the zero value may exhibit real biological variability (18).

### Imputation of dropout values

To impute the gene expression levels, we define hyper-parameter *t* as the threshold to determine *X*_*ij*_ is a dropout event or not. For entry whose dropout probability is less than *t*, we consider it as a real observation, its original value will remain. Otherwise, we conduct the imputation with the aid of information from similar cells utilizing non-negative least squares lasso regression. Please notice that we will not borrow any information from any other dropout events. For *j*-th cell, the objective is:

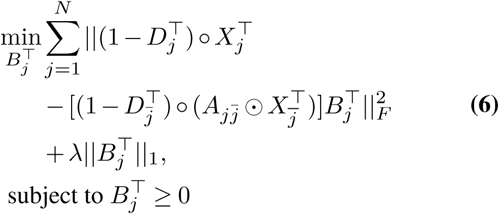

where 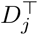 and 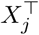 is the *j*-th column of *D* and *X. ○* operator is the Hadamard product which follows (*P ○ Q*)_*ij*_ = *P*_*ij*_*Q*_*ij*_.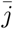 denotes all indices except index *j*, thus 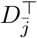 and 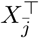 denotes the sub-matrix of *D* and *X* which contains all cells except the *j*-th cell, respectively. 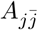 stores the pairwise affinity between *j*-th cell and all other cells; 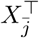 is a sub-matrix of *X* which contains all cells except the *j*-th cell. ⨀ operator represents the vector and matrix multiplication, e.g. (*p* ⨀ *Q*)_*ij*_ = *p*_*i*_*Q*_*ij*_. Leveraging 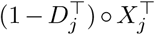 as target indicates that genes with high dropout probability in *j*-th cell will not contribute to optimization. Furthermore, the multiplication of 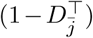 and 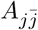 ensures that the information is only borrowed from the trusted genes with low dropout probabilities in the similar cells. Non-negative weights 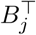 are extra contributions of all other cells learned from regression. Finally, ℒ1 is applied to avoid over-fitting and further ensure the imputation borrow information from the cell’s most similar neighbors.

After obtained the estimated 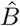, the imputed matrix 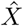 can be calculated. Count values that have dropout probability lower than threshold *t* will remain the same, and values that have dropout probability higher than *t* will be replaced by imputation result.

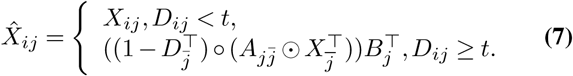

### I-Impute

I-Impute is a “self-consistent” tool to impute scRNA-seq data. As illustrated in **Figure 1B**, it utilizing C-Impute to iteratively refine the SAVER processed data. With *n* times of iterations, the final imputed result remains self-consistent (< 0.1).

We define self-consistency of a functional mapping *f* : *x* → *x* given by input data *X* ∈ *M* × *N* :

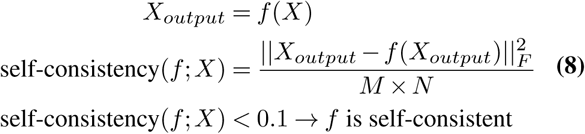

self-consistency(*f* ; *X*) < 0.1 → *f* is self-consistent

## Evaluation metrics

### Adjusted Rand index and normalized mutual information

The adjusted Rand index (ARI) (29) and normalized mutual information (NMI) (30) are adopted as clustering accuracy, which measures the similarity between predicted clustering results and actual clustering labels. A value close to 0 indicates random labelling or no mutual information, and a value of 1 demonstrates 100% accuracy of clustering or perfect correlation.

### Silhouette width

The silhouette width (SW) measures the similarity of a sample to its class compared to other categories (31). It ranges from -1 to 1. A higher silhouette value suggests a more appropriate clustering, a silhouette value near 0 intimates overlapping clusters and a negative value be tokens that the clustering has been performed incorrectly. We adopted the silhouette width to evaluate the model’s imputation power. We used the ground-truth subtype classes as the input cluster labels.

### Simulation and benchmark settings

Splatter are used to generate simulated scRNA-seq data. The parameters used for our simlation dataset are nGroups=3, nGenes=2000, batchCells=150, seeds=42, dropout.type=“experiment”, dropout.shape=-1 and droupout.mid=2, 3, 5 for three different dropout rate data.

SAVER and scImpute are adopted as the competing tools. For SAVER R package, we used the “saver” function with the parameters ncores=12 and estimates.only=TRUE to perform the imputation tasks. The parameters for scImpute are drop_thre=0.5, ncores=10, Kclusters=(number of true clusters in input data).

On synthetic data, I-Impute configuration is n=40, normalize=False, and iteration=True. On real data sets, I-Impute configuration is n=40, and iteration=True for mouse Bladder cell dataset and ES cell dataset and n=20, and iteration=True for mouse Aortic Leukocyte cell dataset.

### Data availability

The real scRNA-seq data used in this study are all publicly available. The mouse ES cell dataset (32) was downloaded from the Gene Expression Omnibus (GEO) with the accession code GSE65525. The mouse Bladder cell dataset and Aortic Leukocyte cell dataset were downloaded from the PanglaoDB (33) with the accession code SRS3044239 and SRS2747908 respectively.

## Code availability

The Python package I-Impute is freely available at https://github.com/xikanfeng2/I-Impute.

## Discussion

In this paper, we introduced I-Impute, which is designed to impute scRNA missing entries iteratively. Experiments using synthetic and real data demonstrated I-Impute to be particularly suited for cell subpopulation discovery.

There are some advantages of I-Impute compared with scImpute and SAVER. Firstly, I-Impute groups SAVER and C-Impute together, and iteratively imputes the missing values until convergence. Adding iteration makes the imputed matrix holds self-consistency and a tighter hierarchical structure. Secondly, scImpute ask the user to decide the cell groups number *k* and assign cells in the same group equal weights during imputation. Here I-Impute gets rid of hyper-parameter *k* and builds a continuous affinity matrix leveraging Gaussian kernel. Last but not least, Lasso regression makes unimportant weights zero, which can help to filter the distant cells for the regression.

The current implementation of I-Impute still has several issues. Firstly, the current regression formula is unable to model the underlying non-linear relationship. We are considering to add deep learning architectures into the I-Impute architecture. Secondly, with the increase of cell and gene numbers, the running time of I-Impute overgrows. We intend to fasten I-Impute running efficiency with the aid of PyTorch tensor parallel computing.

## Conclusions

It is essential to impute the missing values in scRNA-seq before the downstream analysis. We conceived an imputation metric “self-consistency” and proposed an iterative imputation tool, I-Impute. Experiments on simulation data and real data sets established I-Impute feasibility in imputation and discovering the underlying cell subpopulation.

## List of abbreviations

scRNA-seq: Single-cell RNA-sequencing
ARI: Adjusted Rand Index
NMI: Normalized Mutual Information
SW: Silhouette Width
RMSE: Root Mean Square Error

## Ethics approval and consent to participate

Not applicable.

## Consent for publication

Not applicable.

## Acknowledgements

Not applicable.

## Author’s contributions

S. C. L. conceived the idea and supervised the project.

X. F., L. C., Z. W., S. C. L. discussed the algorithm and designed the experiments.

X. F. implemented the code and conducted the analysis.

L. C., X. F. drafted the manuscript.

Z. W., S. C. L. revised the manuscript.

All authors read and approved the final manuscript.

## Supplementary Tables

**Table S1.**
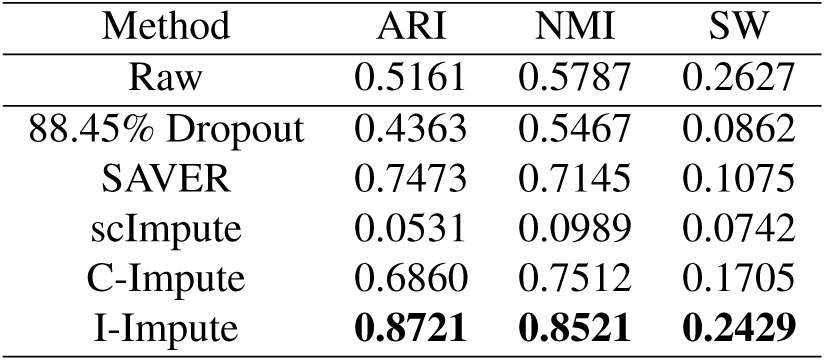
Imputation performance on synthenic data (88.45% dropout)

| Method | ARI | NMI | SW |
| --- | --- | --- | --- |
| Raw | 0.5161 | 0.5787 | 0.2627 |
| 88.45% Dropout | 0.4363 | 0.5467 | 0.0862 |
| SAVER | 0.7473 | 0.7145 | 0.1075 |
| scImpute | 0.0531 | 0.0989 | 0.0742 |
| C-Impute | 0.6860 | 0.7512 | 0.1705 |
| I-Impute | **0.8721** | **0.8521** | **0.2429** |

**Table S2.** Imputation performance on synthenic data (63.29% dropout)

| Method | ARI | NMI | SW |
| --- | --- | --- | --- |
| Raw | 0.5161 | 0.5787 | 0.2627 |
| 63.29% Dropout | 0.3105 | 0.4115 | 0.1910 |
| SAVER | 0.9784 | 0.9700 | 0.2739 |
| scImpute | 0.2033 | 0.3904 | 0.2054 |
| C-Impute | 0.9644 | 0.9509 | 0.2496 |
| I-Impute | **1.0** | **1.0** | **0.3908** |

**Table S3.** Imputation performance on synthenic data (45.16% dropout)

| Method | ARI | NMI | SW |
| --- | --- | --- | --- |
| Raw | 0.5161 | 0.5787 | 0.2627 |
| 45.16% Dropout | 0.6000 | 0.6306 | 0.2137 |
| SAVER | 0.9801 | 0.9710 | 0.3292 |
| scImpute | 0.3596 | 0.4962 | 0.2365 |
| C-Impute | 0.4727 | 0.5551 | 0.2566 |
| I-Impute | **0.9801** | **0.9710** | **0.4123** |

**Table S4.** Pearson correlation result on synthenic data.

| Method | 88.45% Dropout | 63.29% Dropout | 45.16% Dropout |
| --- | --- | --- | --- |
| SAVER | 0.5836 | 0.7915 | 0.8764 |
| scImpute | 0.6451 | 0.8582 | 0.9263 |
| C-Impute | 0.7611 | 0.8873 | 0.9305 |
| I-Impute | **0.7812** | **0.8993** | **0.9357** |

**Table S5.** Imputation performance on mouse Bladder cells data.

| Method | ARI | NMI | SW |
| --- | --- | --- | --- |
| Raw | 0.1937 | 0.4500 | 0.0737 |
| SAVER | 0.5253 | 0.7085 | 0.0621 |
| scImpute | 0.1937 | 0.4500 | 0.0686 |
| C-Impute | 0.1664 | 0.4317 | 0.0741 |
| I-Impute | **0.6054** | **0.7892** | **0.1758** |

**Table S6.** Imputation performance on mouse ES cells data.

| Method | ARI | NMI | SW |
| --- | --- | --- | --- |
| Raw | 0.2410 | 0.5160 | 0.0353 |
| SAVER | 0.6920 | 0.7329 | 0.2235 |
| scImpute | 0.3574 | 0.5258 | 0.0418 |
| C-Impute | 0.2410 | 0.5160 | 0.0411 |
| I-Impute | **0.7047** | **0.7444** | **0.2275** |

**Table S7.** Imputation performance on mouse Aortic Leukocyte cells data.

| Method | ARI | NMI | SW |
| --- | --- | --- | --- |
| Raw | 0.3463 | 0.6352 | 0.0290 |
| SAVER | 0.3605 | 0.6837 | **0.1075** |
| scImpute | 0.3377 | 0.6237 | 0.0358 |
| C-Impute | 0.3427 | 0.6398 | 0.0206 |
| I-Impute | **0.5220** | **0.7728** | 0.0711 |

